# Neurothreads: Cryogel carrier-based differentiation and delivery of mature neurons in the treatment of Parkinson’s disease

**DOI:** 10.1101/2020.01.30.927244

**Authors:** Aleksandra Filippova, Fabien Bonini, Liudmila Efremova, Olivier Preynat-Seauve, Amélie Béduer, Karl-Heinz Krause, Thomas Braschler

## Abstract

We present *in-vivo* transplantation of mature dopaminergic neurons by means of macroporous, injectable carriers, to enhance cell therapy in Parkinson’s disease. The carriers are synthesized by crosslinking carboxymethylcellulose at subzero temperatures, resulting in cylindrical, highly resilient porous cryogels, which we term Neurothreads. We develop efficient covalent immobilization of the neural adhesion proteins laminin 111, collagen IV and fibronectin, as well as of the extracellular matrix extract Matrigel to the Neurothreads. We observe the highest neural spreading on laminin 111 and Matrigel. We show compatibility with established dopaminergic differentiation of both HS420 human embryonic stem cells and the LUHMES midbrain model cell line. The porous Neurothread carriers withstand compression during minimally invasive stereotactic injection, and ensure viability of mature neurons including extended neurites. Implanted into the striatum in mice, the Neurothreads enable survival of transplanted mature neurons obtained by directed differentiation of the HS420 human embryonic stem cells, as a dense tissue *in situ*, including dopaminergic cells. With the successful *in-vivo* transfer of intact, mature and fully open 3D neural networks, we provide a powerful tool to extend established differentiation protocols to higher maturity and to enhance preconfigured neural network transplantation.

## Introduction

Parkinson’s disease is a major neurodegenerative disorder currently affecting around ten million people worldwide^1^. There are various symptomatic treatments ranging from oral medications to implantation of stimulation electrodes ^2,3^. At present, there is no widely applicable cure and symptomatic treatment success is highly variable, depending on the disease stage, individual factors, treatment modalities and other, some yet unknown factors^2,4^. On the cellular level, Parkinson’s disease is characterized by progressive degeneration of dopaminergic neurons within *substantia nigra pars compacta* (SNc) and their projections, largely towards the striatum – the area involved in movement control^2^. The decrease in dopamine levels within the striatum, in its turn, leads to impairment of motor functions and development of symptoms, such as rigidity, tremors and bradykinesia^2^.

Fetal mesencephalic tissue transplantation has shown promising results in the treatment of Parkinson’s disease patients in clinical trials^4,5^. Currently, more ethical and practical cell replacement approaches are being pursued, with the aim to ease localization of immune-matching cell sources and to simplify and standardize graft preparation^6^. Numerous pre-clinical^7–13^ and emerging clinical^9,14,15^ stem cell studies are investigating intrastriatal transplantation of neural precursor cells (NPCs), derived from autologous or amplifiable human sources such as human embryonic stem cells (hESCs)^7–9^, induced pluripotent stem cells (iPSCs)^7,9,10^, parthenogenetic stem cells^15,16^ or direct reprogramming of autologous somatic cells^11–13^.

To provide dopaminergic neuron replenishment in Parkinson’s disease^2^, stem-cell derived, dopaminergically primed NPCs have been shown to possess acceptable survival rates *in vivo* and to improve motor function in animal models of the disease^4,9,16^. However, transplantation of immature cells implies lack of control over the maturation stage, risking variability in mature phenotype of the graft and thus variability in clinical outcome^2,4^. Additionally, there is a risk of debilitating or life-threatening side-effects, such as tumorigenesis - due to remaining proliferating cells - as well as graft-induced dyskinesias (GIDs) of unknown origin, previously observed in some patients with fetal transplants^2,4^. The differentiation and malignancy potential of a given iPSC line remains difficult to predict^17^, and pre-transplantation quality assessment at later time points can be expected to reveal more pertinent information regarding the *in-vivo* outcome. Mature neurons however have an overly complex and fragile architecture, and thus are poorly tolerant to pre-transplantational handling procedures, particularly enzymatic dissociation^4,18^. This raises the challenge of how to transplant mature neurons while protecting their fragile architecture. A solution to this would enable more mature cell delivery in Parkinson’s disease, and more generally transplantation of 3D preconfigured neural circuitry, truly feasible at present feasible only with embryonic transplants.^19^

Structural support in the form of biomaterials has been reported to enhance the survival rate of more or less well differentiated neuronal cells during handling and transplantation^18,20,21^. Encapsulating biomaterials have for example been used to enhance transplantation of immature neural progenitor cells, including embryonic primary ventral midbrain cells^22,23^, into the rodent brain as a potential treatment for stroke^24^ and Parkinson’s disease^18,22,23^. A further promising approach for more strongly anchorage-dependent cells is the use of a suspension of microscaffold carriers.^21,25,26^ While improving aspects of cell survival, these studies do not address the question of how a fully organized and mature neural network might be transplanted. Nerve-guide inspired cylindrical sheaths have been shown to successfully maintain mature tissue integrity^27–29^. While offering the exciting possibility to transplant neural tracts intra-cerebrally in small animal models^28,29^, by construction, these implants physically limit lateral access for vascularization required for larger volumes^30^.

To provide true 3D transplantation, the use of slowly biodegradable highly porous biomaterial scaffolds has been proposed^20,21^: such scaffolds can be used for robust culture of primary neurons and neuron-like cells *in vitro*, and have been shown to enhance survival during *in-vitro* handling procedures^20,21^. To facilitate minimally invasive cell delivery *in vivo*, cryogels as a class of particularly rugged macroporous hydrogels have been advanced^26,31,32^. Cryogels are synthesized by freezing a hydrogel premix, following by polymerization in the cold^33^. Upon thawing, highly resilient sponge-like structures with macropores on the order of tens to hundreds of micrometers are obtained.^32,34^ The extreme porosity allows the desired favourable interaction with the surrounding tissue, including colonization and vascularization of the pore space^26,34^ and facile exchange with the surrounding tissue^35^. A series of cryogels, including gelatin-laminin, heparin, alginate and CMC-based cryogels were previously demonstrated to be compatible with neuronal culture^20,21,36^. Some of these materials have been shown to withstand delivery through fine needles^20,31^, thereby providing the potential for stereotactic minimally invasive grafting procedures.

Our aim is indeed to implement macroporous cryogel scaffolds^20,37^ to enable minimally invasive, stereotactic implantation of mature dopaminergic neurons *in-vivo*. For neural progenitor cell seeding and *in-situ* maturation, we require an injectable and biocompatible scaffold with stably bound neural adhesion molecules. The gels should further be compatible with established dopaminergic differentiation protocols^38^. Our goal is to address the challenge of *in-vivo* injectability of mature neurons, ultimately providing a later and safer grafting option for treatment of Parkinson’s disease and beyond.

## Materials and methods

We aim at obtaining *in-vivo* injectability of mature neurons by the mechanical support of tailored cylindrical cryogels, which we term “Neurothreads” (Fig. 1). To achieve this aim, we synthesize and functionalize cryogels (Fig. 1a), develop and assess cell seeding and on-gel differentiation and maturation methods (Fig. 1b, 1c), develop and characterize steoreotactic injection and finally assess neural survival and phenotype in a pilot *in-vivo* (mouse) model (Fig. 1d).

**Fig 1.**
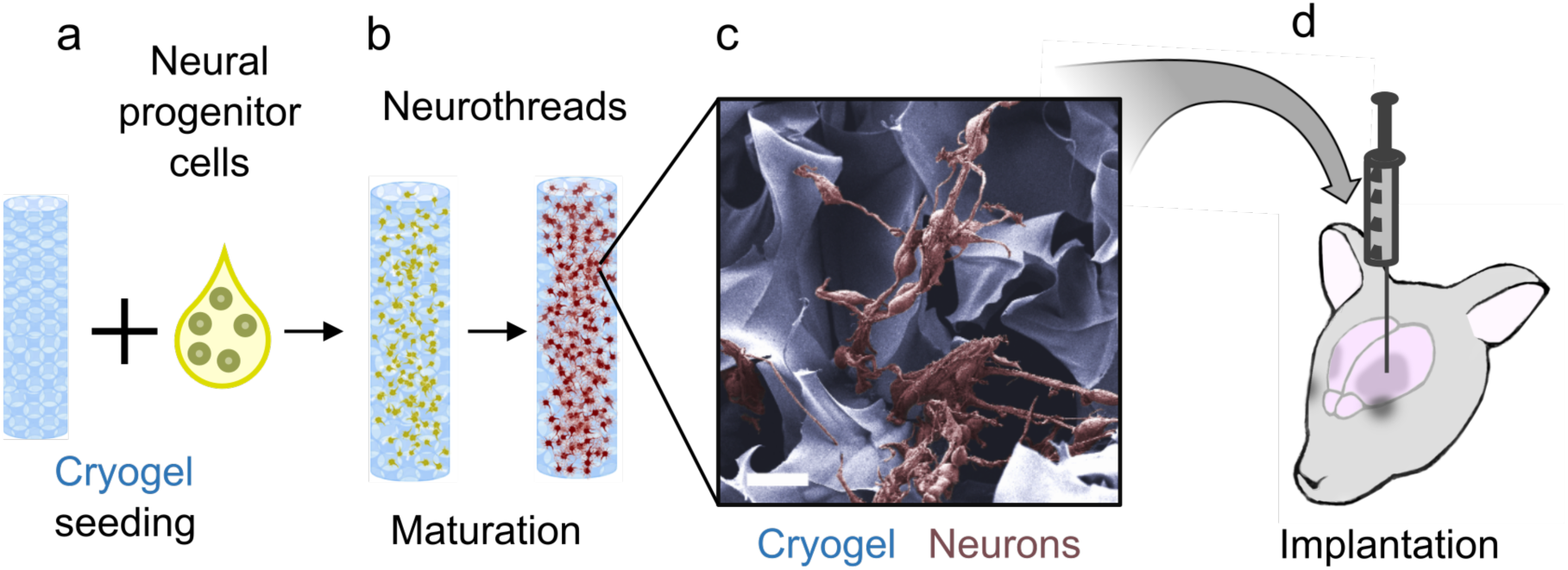
Cryogel carriers as support of mature neurons during transplantation. a) For transplantation of mature neurons, neural progenitor cells are seeded on porous cylindrical cryogels termed Neurothreads. b) The cells on the Neurothreads are then matured in-situ, to c) form an extensive neural network supported by the Neurothread cryogel carriers. d) The ultimate aim is to transplant intact mature neural networks, for dopamine replenishment in Parkinson’s disease. Here, we assess neural survival in a murine model. c) is a manually colored scanning electron microscope (SEM) image of LUHMES^39^-derived neurons, matured on a laminin 111-coated CMC cryogel. Scale bar = 20 µm.

### Cryogel preparation

Cryogel mix was prepared, as described previously^20^ with minor changes. Briefly, 1.5% carboxymethylcellulose (Hänseler, Carmellose sodium 500) and 0.05 % adipic dihydrazide (Sigma) were dissolved in 50 mM PIPES (Sigma-Aldrich) buffer. The pH was adjusted to 6.7 with sodium hydroxide (Sigma). For induction of cryogel synthesis, the water-soluble carbodiimide N-(3-dimethylaminopropyl)-N-ethylcarbodiimide hydrochloride (EDC, Sigma) was added to the premix, which was then moulded into a desired shape and put to freeze - maximum 3 minutes after EDC addition. In detail, for Neurothread microcylinder fabrication, the mix was injected into glass microtubes with inner diameters of 0.58mm (hitherto referred to as 0.6mm), 0.4 mm and 0.3 mm (World Precision Instruments) and placed at -10°C for 48 hours. To increase the available cell amount for histological and RT-PCR analysis, we also fabricated larger disk-shaped cryogels. For this, the reaction mix was poured between two glass slides, separated by 1 mm high spacers and placed at -20°C for 48 hours.

Upon thawing, the disc gels were stamped into disc shapes with a diameter of 4 mm. Then, they were washed on Stericups (Merck Millipore), using dehydration/rehydration cycles with sterile de-ionized (DI) water, followed by 1M sodium hydroxyde incubation for 1 hour, 10 mM pH 8 ethylene-diamine-tetraacetic acid (EDTA, Sigma-Aldrich) for 15 minutes, washed with deionized (DI) water and phosphate buffered saline PBS, and finally, autoclaved at 121°C for 20 minutes in PBS. Similar steps were used for washing the Neurothreads, with the exception that they were vortexed in washing solutions, instead of being dehydrated using Stericups, to prevent mechanical damage.

### Cryogel functionalization

The cryogels were functionalized in sterile conditions, using one of the following extracellular matrix (ECM) protein products: laminin (Laminin 111, Sigma L2020, from mouse Engelbreth-Holm-Swarm sarcoma), fibronectin (Sigma F1141, from bovine plasma), collagen IV (Sigma C5533, from human placenta), or Matrigel (Sigma, E1270). For fluid exchange during functionalization, vacuum pressure adjusted to 3 kPa was used for dehydration/rehydration of the disc cryogels on Stericups. Neurothread cryogel carriers were functionalized by vortexing with coating solutions.

The detailed protocol was as follows: Firstly, the gels were pre-equilibrated to a desired pH with 100 mM of pH buffer (acetic acid for pH 4 and 5, MES for pH 6 and 7 and HEPES for pH 8). Afterwards, they were incubated for 1h at room temperature or overnight at +4°C with a coating solution, consisting of 100 mM buffer, 1M urea and 250 µg/ml coating protein for functionalization adjustment experiments and 500 µg/ml laminin for the rest of the experiments. For negative controls, de-ionized water, instead of proteins, was added into the coating solution at pH 6, while the rest of the components remained the same. In the case of Matrigel coating, gels were pre-equilibrated with pH 6 buffer and incubated with a solution, consisting of Matrigel, diluted 1:50 in Advanced DMEM/F12 medium (Thermofisher Gibco brand, cat # 12634010) and 1M Urea (final Matrigel protein concentration 200 µg/ml). In order to covalently crosslink the coating molecules to the cryogel, after a washing step with DI water, the gels were incubated with EDC (diluted in 0.5 M pH 5.5 MES buffer), followed by 25 mM pH 10 bicarbonate buffer (Sigma) and once again washed with DI or PBS. Coated gels were stored at +4°C in PBS for a maximum of one week.

### Evaluation of the functionalization efficiency

For quantification of protein binding, the bound protein was labelled by isothiocyanate chemistry^40^. Specifically, coated or control cryogels were incubated with 350 µg/ml Rhodamine B isothiocyanate (RITC, Sigma), dissolved in 70 mM pH 10 bicarbonate buffer with 20% isopropanol. The cryogels were incubated with the labelling solution overnight at 4°C. The next day, unreacted dye was removed by washing the gels multiple times with a 0.1 M pH 9.3 bicarbonate buffer containing 20% isopropanol, then the same buffer without isopropanol, and finally DI water. To enable quantification of the bound dye, each purified gel was dissolved in 100 µl 1 M NaOH by autoclaving at 120°C for 20 minutes. Fluorescence intensity of the obtained solution was measured quickly after the dissolution of the gels, to prevent aggregate formation, on a Spectramax Paradigm plate reader (Molecular Devices) with 540 nm excitation and 580 nm emission wavelengths. The amount of bound proteins was estimated from the detected amount of RITC by assuming a 50% labelling efficiency on the available aminogroups deduced from the fraction of lysine amino acid residues in the primary sequences (neXtProt^41^ sequences NX_P25391, NX_P07942 and NX_P11047 for the three subunits of laminin111, NX_P11047 for fibronectin, and NX_P02462, NX_P08572, NX_Q01955, NX_P53420, NX_P29400, NX_Q14031 for the 6 possible subunits of collagen IV). This gives a conservative estimate of the protein amount, as the actual labelling efficiency is known to show saturation at higher labelling reagent concentrations^42^. Finally, by comparison to the known amount of protein provided during the coating procedure per gel, the reaction efficiency was determined.

### Cell culture and differentiation

The LUHMES cell line^39^ (ATCC CRL-2927) was kindly provided by Prof. Marcel Leist, University of Konstanz. The cells were maintained and passaged by established protocols^43^. T75 flasks were coated with 0.0015 % poly-L-ornithine (PLO) (EDM Millipore) and 1 µg/ml fibronectin (Sigma-Aldrich). During maintenance, the cells were passaged every two days and kept in Advanced DMEM/F12 medium (Gibco 12634010), supplemented with 1:100 N2 supplement (Gibco), 100 ng/ml FGF2 (Cell Guidance systems), 2 mM L-Glutamine (Gibco) and 1:100 penicillin-streptomycin (PS) (Gibco).

Differentiation medium (DM) consisted of Advanced DMEM/F12, supplemented with N2 supplement, 2 mM L-glutamine, 100 mM db-cAMP (Sigma-Aldrich), 1 mg/ml Tetracycline (Sigma-Aldrich) and 20 ng/ml GDNF (Cell Guidance Systems). For differentiation^43^, LUHMES cells were passed into a new T75 culture flask (ThermoScientific) on the day before starting the differentiation. On day 0, the medium was changed for DM. After 48 hours of pre-differentiation, cells were seeded either onto well plates, coated with 0.0015 % PLO and 1 µg/ml laminin 111 or on variously functionalized cryogels at a density of 100’000 cells per gel for the larger disk gels and 30’000 cells per gel for the smaller Neurothread microcylinder gels. The LUHMES cells were then further differentiated in DM until day 7 (D7) prior to further analysis.

The HS420 hESC cell line was kindly provided by Outi Hovatta, Karolinska Institutet^44^. Cells were grown on T75 flasks, coated with 0.5 µg/cm^2^ rhLaminin-521 (Biolamina LN521), diluted in PBS +Ca^2+^/ +Mg^2+^ (Gibco). StemFlex medium (Gibco, A3349401) was used for cell maintenance. During culture, medium was changed every 2 days and cells were passaged roughly every 4 days.

hESCs were differentiated according to a published protocol^38^. For differentiation, hESCs were first seeded at a density of 10 000 cells/cm^2^ into a 6 well plate (Corning), coated with 1 µg/cm^2^ rhLaminin-521, diluted in PBS +Ca^2+^/ +Mg^2+^. The cells were differentiated for 8 days in medium I, consisting of 50% DMEM/F-12 with HEPES and L-glutamine (Gibco 31330038), 50% Neurobasal (Gibco 21103049), 1:100 N2 supplement, 1:100 B27 supplement minus vitamin A (Gibco), 1 mM L-glutamine, 1% vol/vol penicillin-streptomycine (PS, Gibco) with addition of 10 µM SB431542 (Sigma-Aldrich, 301836-41-9), 100 ng/ml Noggin (Miltenyi Biotech), 300 ng/ml SHH-C24II (Miltenyi Biotech), and 1 µM CHIR99021 (Cell Guidance Systems). Additionally, 10 µM Y-27632 (Cell Guidance Systems) was used for the first two days after seeding to enhance cell survival. The medium was changed every two days. On day 9, the medium was changed to medium I, with addition of 100 ng/ml FGF8b (Cell Guidance Systems). On day 11, the cells were passaged at density of 500 000 cells/cm^2^ into a new PLO/rh laminin-521 coated 6 well plate. Medium II consisted of Neurobasal, 1:50 B27 supplement without vitamin A, 2mM L-glutamine, 1% vol/vol penicillin-streptomycine, supplemented with 100 ng/ml FGF8b, 20 ng/ml BDNF (Cell Guidance systems) and 0.2 mM L-ascorbic acid (AA) (Sigma-Aldrich). It was used for differentiation between day 11 and day 16 or 17 depending on the experimental run. Additionally 10 µM Y-27632 was added in the medium for the first two days after seeding. The medium was changed every two days or every day, if it turned yellow. On day 16 or 17, pre-differentiated cells were dissociated and seeded onto the cryogels at a density of 30 000 cells per Neurothread cryogel respectively 60’000 cells per disc cryogel. Cryogels were placed into a 6 well plate at 4-5 gels/well (disk gels) or at 10 gels/well (Neurothreads) and differentiated until day 40 in medium III, consisting of Neurobasal, 1:50 B27 supplement without vitamin A, 2mM L-glutamine, 1% vol/vol penicillin-streptomycine, 20 ng/ml BDNF, 0.2 mM AA, 10 ng/ml GDNF, 500 µM db-cAMP, 1 µM DAPT (Cell Guidance Systems). The medium was changed every 2-3 days until day 40 (D40), when the samples were collected for analysis. Quantification of the total viable cell number by Alamar Blue reduction was carried out on a minority of the gels 24h post seeding (referred to as “D17”), and on day 40 (referred to as “D40”).

### Immunocytochemistry, immunohistochemistry, and imaging of in-vitro samples

Cryogels or well plates, containing LUHMES cells and hESCs, were fixed using 2% para-formaldehyde (PFA), then incubated with 0.5 % triton-X and blocking buffer, containing 1% fetal bovine serum (FBS) (Gibco) and 0.5% triton-X in PBS. Primary antibodies, diluted in PBS with 1% FBS and 0.5% triton-X were added to the samples and left for incubation at 4°C overnight on a waving rotator. The next day, primary antibodies were replaced by the secondary antibodies and incubated for 3 hours on a rocking platform. Afterwards, samples were incubated with 4’,6-diamidin-2-phenylindole DAPI, washed and imaged. All antibodies used here are specified in Supplementary 4 (Table S4). Imaging of the cryogels was done on a Zeiss LSM 800 confocal microscope.

For differentiation analysis, cryogels were cut into smaller pieces and confocal z-stacks of 15 µm height were taken at 20x magnification within the thickness of the cryogel. The stacks were projected down and the marker-positive cells were quantified manually. All 2D plate-cultured cells were imaged using a plate reader ImageXpress XL (Molecular Devices) or microscope Nikon Eclipse Ts2 directly on the plate.

### Quantification of neural spreading

Quantification of neural spreading was carried out on disk-shaped cryogels functionalized with laminin 111, fibronectin and collagen IV, as well as positive (Matrigel) and negative (no ECM protein) controls. LUHMES cells were seeded and differentiated on the gels as described above, followed by fixation and labelling with anti BIII-tubulin (TUBB) – a neural cytoskeleton marker - and DAPI and imaged as a z-stack on a confocal microscope. For quantification of neural spreading, a 50 µm z-stack of cryogels was taken from the top of each gel and projected down, using ImageJ software. The network score was calculated by:

1. subtracting DAPI-positive area from BIII-tubulin images (to quantify only the neurites and not the cell bodies),
2. quantifying the number of neurites as the remaining BIII-tubulin positive objects,
3. Normalizing the number of neurites to the number of DAPI-labeled nuclei.

In summary: 

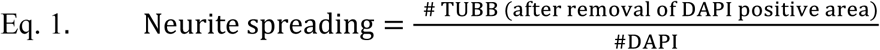

### Immunohistochemistry of in-vivo samples

Immunohistochemistry of *in-vivo* grafts was done according to established procedures^45^: Murine brains embedded into paraffin blocks were cut into 7µm coronal sections, mounted on Superfrost™ glass slides (ThermoFisher), and dried overnight at 37°C. The sections were hydrated by submergence into xylol, followed by decreasing ethanol gradient (100% ethanol, 95%:5% ethanol:DI water, 70%:30% ethanol:DI water) baths. Epitope retrieval was then done using pH 6 0.01 M citrate buffer, for 15 minutes at 95°C. The slices were then blocked with an alkaline phosphatase (AP) blocking solution (BLOXALL, Vector labs) for 10 minutes, and additionally for one hour with 30 µg/ml murine Fab fragments in 0.2% triton-X PBS solution to further decrease unspecific staining due to the murine primary antibody, and finally with with 10% fetal bovine serum (FBS) in 0.2% triton PBS solution for 30 minutes. The first primary antibody (STEM121), diluted in 1% horse serum 0.2% triton PBS solution was added onto the slides and incubated for one hour. It was followed by incubation with an AP-conjugated IgG protein for 30 minutes (Vector labs) and developed with an AP substrate (Vector Red, Vector labs). Next, slices were incubated with the remainder of the primary antibodies (TH, Neurofilament 200), diluted in 1% FBS for one hour, followed by an hour incubation with secondary fluorescent antibodies and, lastly, DAPI. The slices were mounted using Fluorsave (Sigma) and imaged using a widefield microscope with structural illumination (ZEISS Apotome 2). Further details on the antibodies used can be found in Supplementary 4 (Table S4).

For hematoxyline/eosine (HE) staining, paraffine-embedded rehydrated slices were submerged into hematoxylin (Merck) for 5 minutes, flushed with DI water for 2-3 minutes, submerged into eosine Y (Sigma) for 5 minutes, rapidly rinsed with DI water, de-hydrated, using a rising gradient of ethanol (70%, 95%, 100%), followed by Neo-Clear (Merck) and fixed using mounting medium Neo-Mount (Merck). Imaging was performed with Nikon Eclipse Ts2.

### Viability assays

Evaluation of the total number of viable cells within the cryogels was done using Alamar Blue assay (Invitrogen) according to the manufacturer’s instructions. A standard curve with a known number of pre-differentiated LUHMES cells or HS420-derived NPCs was used. Alamar blue reduction as a measure of cell metabolism was measured at 550 nm excitation and 590 nm emission wavelengths on Spectramax Paradigm.

Live/dead staining was achieved by incubation of cryogels and well-plate cultured cells in culture medium, containing 3 µM Calcein AM (Sigma) for 1 hour and then adding 3 µM propidium iodide (PI) (Sigma) for additional 40 minutes. The cells in 2D cultures were imaged with Nikon Eclipse Ts2, without fixation. The cryogels were fixed in 2 % PFA after the live/dead staining, cut into thin vertical slices and imaged using Zeiss LSM 800. Live and dead cells were counted manually; statistical analysis was carried out on the recovery of the life cells as the primary readout, since we anticipate dead cells to be more easily fragmented or lost.

### Gene expression analysis

RNA was extracted from samples using RNeasy micro kit (QIAGEN) according to the manufacturer’s instructions. For cryogels, up to 5 disk-shaped gels from a given well were pooled prior to extraction to obtain sufficient RNA (250ng per reaction, Nanodrop 2000c, ThermoScientific). cDNA was synthesized using a mix of random hexamers – oligo d(T) primers and PrimerScript reverse transcriptase enzyme (Takara bio inc. Kit), again following the manufacturer’s instructions. SYBR green assays were designed using the program Primer Express v 2.0 (Applied Biosystems) with default parameters. Primers were obtained from Microsynth AG - sequences are specified in Supplementary 5 (Table S5). PCR reactions (10 μl volume) contained diluted cDNA, 2 x Power SYBR Green Master Mix (Applied Biosystems), 300 nM of forward and reverse primers. PCR were performed on a SDS 7900 HT instrument (Applied Biosystems) with the following parameters: 50°C for two minutes, 95°C for ten minutes, and 45 cycles of 95°C 15 secondes-60°C one minute. Each reaction was performed in three replicates on 384-well plate. Raw Ct values obtained with SDS 2.2 (Applied Biosystems) were imported in Excel. The Ct values were converted to gene expression levels by assuming full doubling efficacy per cycle. For evaluation of relative changes in gene expression upon differentiation, the expression level was normalized to the maximum expression level for each gene, followed by a second normalization step to the house-keeping gene expression level: GAPDH for the Luhmes differentiation experiments, geometric mean of GAPDH and EEF1A1 according to the geNorm^46^ method for the hESC differentiation experiments. For wells not having reached the threshold by the end of the PCR reaction, evaluation was carried out with a maximum Ct value of 40, which is slightly above the maximum observed in positive reactions, instead.

### Neurothread cryogel handling procedures

Neurothread cryogel carriers with a cylindrical shape of 5 mm height and 0.3mm, 0.4mm and 0.6mm diameter were seeded with pre-differentiated LUHMES cells and cultured under differentiation conditions until day 7. Sterile 24 GA syringes (Hamilton, 0.311mm inner needle diameter according to the supplier’s notice) were used for all injection experiments. For viability testing, cryogels were kept inside the Hamilton syringe needle for one minute and re-injected into new well plates with differentiation medium. 24 hours later, live/dead stain was applied, as described previously. Cryogels were fixed with 2% PFA and analysed using Zeiss LSM 800 confocal microscope. Cells differentiated on laminin/PLO-coated well plates were used as control. They were dissociated from the wells with 0.05% trypsin-EDTA (Gibco), dissolved, centrifuged, re-suspended with DM and (after 1 min in the Hamilton syringe needle) injected into well plates (coated with PLO/laminin). 24 hours later, live/dead stain was applied and, after incubation, wells were imaged without fixation using a Nikon Eclipse Ts2.

For testing injection into soft matter, we fabricated a brain phantom^47^ consisting of 0.2 % agar, dissolved in sterile DI water by autoclaving. The brain phantom was pre-incubated in LUHMES differentiation medium overnight before the experiments.

### In vivo implantation and brain sample collection

All animal experiments were carried out as approved by the Geneva Local Veterinary Committee (Authorization GE/191/19) and in accordance with relevant Swiss regulations. Mice were housed in the specific-pathogen-free (SPF) area of the local animal facility at controlled temperature (22±2°C), with 12h dark/light cycles. Water and food was provided *ad libitum*.

NOD/SCID J mice (Charles River) were injected with D40 hESC-derived neurons seeded on 0.4 mm diameter Neurothread cryogels. For this, we used a 24 GA Hamilton syringe, placed on a stereotactic frame. Opioid analgesic (Buprenophine, 0.1 µg/g weight) was injected intraperitoneally 30min prior to the surgeries and local anaesthesia (Lidocaine) was applied to the scalp prior to incision. Injections were performed under general anaesthesia (Isoflurane). The following coordinates were used for injection: AP: 0.6, ML: 2, DV: 3-2.5 from dura. Implantation of the graft was done synchronously with the slow retraction of the needle. Saline control was injected into the second hemisphere (AP: 0.6, ML: -2, DV: 3-2.5) in a similar manner. Four weeks post-injection, the mice were euthanized with 0.15 mg/g weight pentobarbital and perfused intracardiacally - first with 0.9% NaCl, then with 4% PFA. The brains were collected, dehydrated – using a rising gradient of ethanol and xylol - and embedded in paraffin. Seven µm coronal sections of striatum were obtained using a microtome (Microtome HM 335 E, GMI) and treated for immunohistochemistry or histological analysis.

### Statistical methods

For physical, chemical and cell viability measurements (Figures 2b, 2c, 5b, 6c) n=3-11 and more typically n=3-5 replicates were used, as indicated. For measurements involving cell morphology and differentiation (Figures 3c-3e, 4b, 4c, 6b), N=3-5 independent experiments with 1-5 replicates per experiment and condition were used (overall *n* per condition of 8-18, and more typically 10-12). In the case of RT-PCR on cryogels, each sample further consisted of physically pooled RNA of typically 4-5 cryogels to provide sufficient RNA. To avoid inflation of P-values due to the clustered structure of the data^48^, either the replicate values per experiment and condition were averaged prior to comparison of conditions and statistical analysis carried out using the average results per independent experiment (Fig. 3c, 3d, 4b, 4c, 6b), or relevant stratification covariates were explicitly included into regression analysis (Fig. 3e).

**Figure 2.**
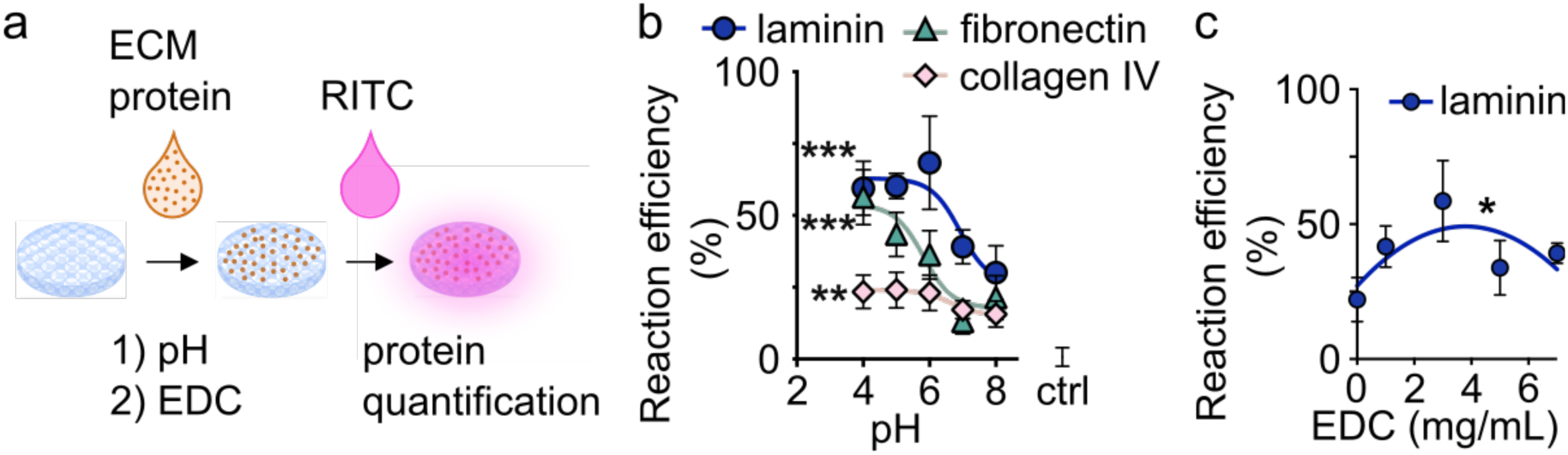
Optimization of CMC cryogel functionalization. a) The coating procedure consists of two main steps: 1) Extracellular matrix (ECM) protein adsorption to the CMC cryogel biomaterial at a controlled pH, followed by 2) covalent immobilization by the action of the water soluble carbodiimide EDC. The amount of bound protein is quantified by the uptake of the amino-reactive dye rhodamine-isothiocyanate^40^ RITC by the covalently modified or control scaffolds. b) The pH applied during the ECM adsorption protein adsorption step, as well as the characteristics of the ECM protein have a profound impact on the immobilization reaction efficiency. c) Reaction efficiency, as quantified by RITC uptake, shows a non-linear dependency on the crosslinker concentration, with an optimum in the lower mg/mL range. n=5-6 per condition, F-tests based on the proportion of variance explained by the non-linear fits in b) and c). *p ≤ 0.05, **p ≤ 0.01, ***p ≤ 0.001, otherwise not significant.

**Figure 3.**
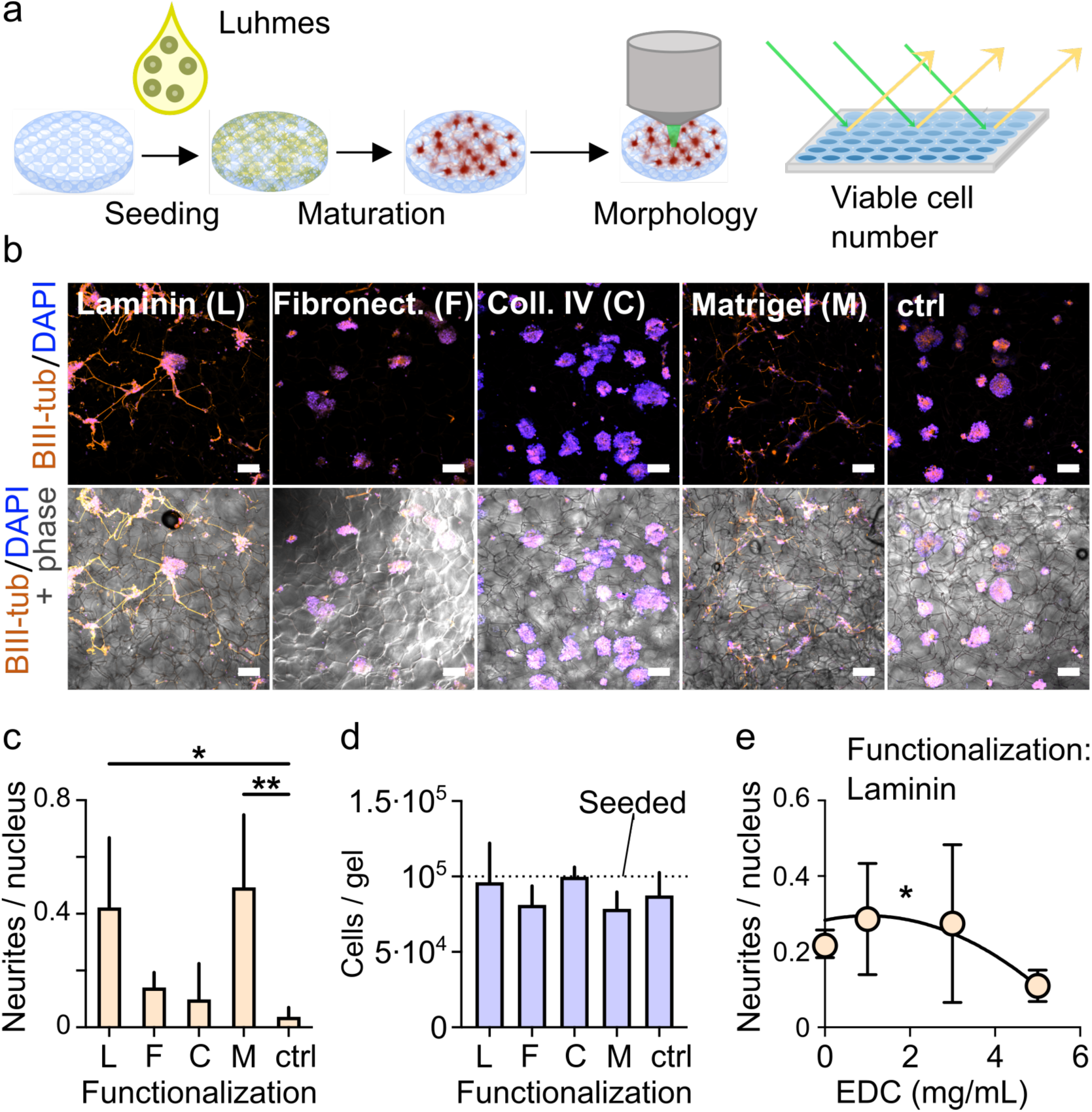
Evaluation of neural adhesion to the functionalized cryogel scaffolds. a) Variously coated cryogel scaffolds were seeded with the LUHMES model cell line. The LUHMES cells were then differentiated in-situ for 7 days by established protocols,^43^ followed by evaluation of neural adhesion by morphology and viable cell presence. b) Confocal images of cryogels stained for BIII-tubulin (neural cytoskeleton marker) and DNA (nuclei, DAPI). Neural adhesion is revealed by neurite sprouting on cryogels coated with laminin 111 (L) and Matrigel (M, positive control), but not on the ones functionalized with fibronectin (F), collagen IV (C), or without functionalization (ctrl, negative control). c) Neural adhesion scoring based on the ratio of neurite to nuclei counts. d) Number of live cells in the variously functionalized cryogels quantified by the Alamar Blue reduction assay. e) Neural adhesion as a function of crosslinker (EDC) concentration used, scoring using the ratio of neurite to nuclear counts. Scale bars in b) represent 100 µm. N=3-5 independent experimental runs, with 2-5 cryogels evaluated for each run and condition (total n=9-18). Statistical analysis on averages per experiment in c) and d), using one-way ANOVA with Dunnett’s correction to compare all conditions with control conditions. Multivariate linear regression in d), using a quadratic term in EDC, and the DAPI count as a covariate. *p ≤ 0.05, **p ≤ 0.01, otherwise not significant.

**Figure 4.**
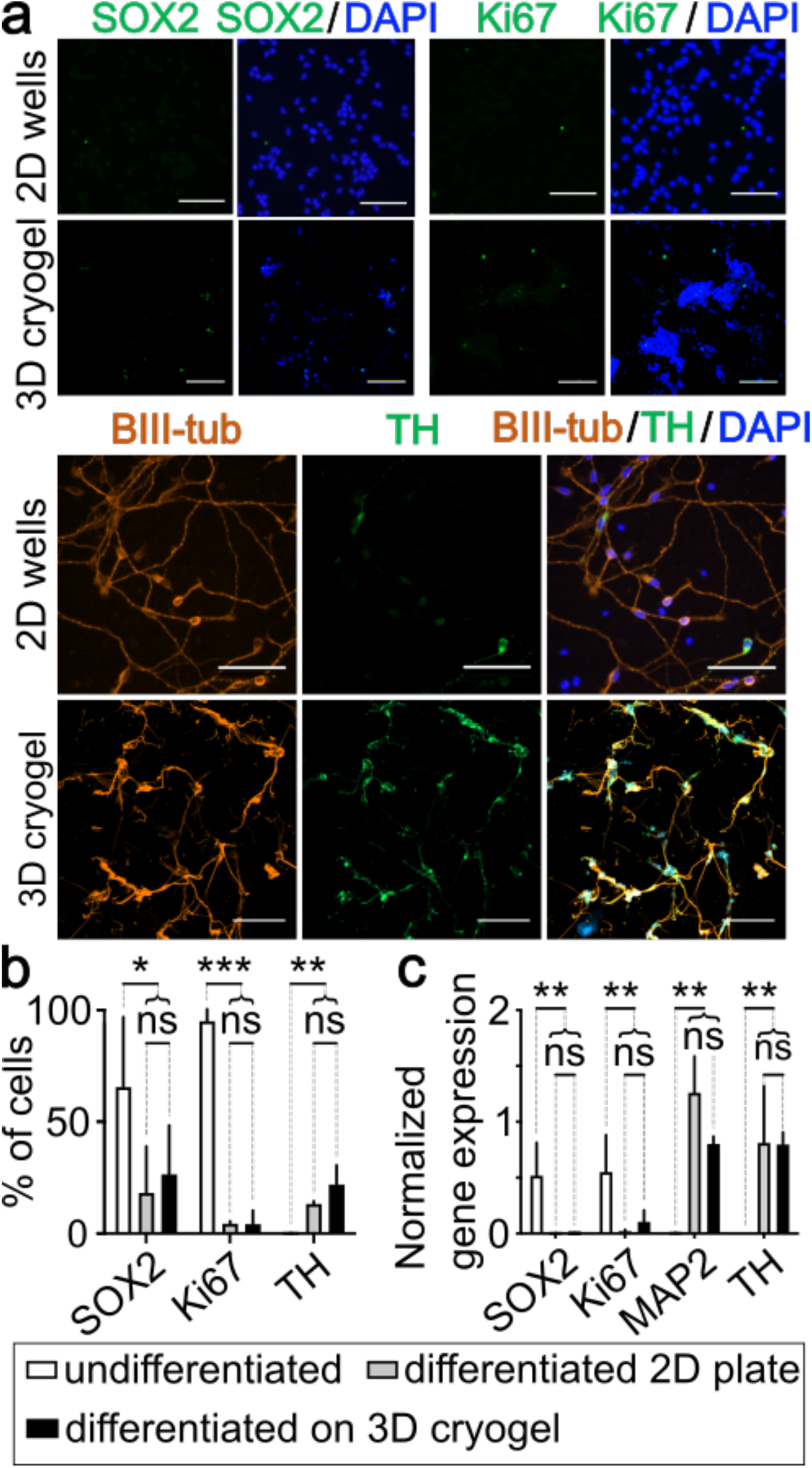
Comparison of differentiation quality between culture plate (2D) and 3D cryogel-cultured cells in LUHMES cells. a) Immunocytochemistry for Ki67 (proliferation marker), SOX2 (pluripotency marker), BIII tubulin (neural cytoskeleton marker) and TH (dopaminergic differentiation marker tyrosin hydroxylase) in differentiated cells on 2D culture wells or inside cryogels. The scale bars are 100 µm; cryogel images are projections of 50 µm z-stacks. b) Quantitative comparison of the fraction of cells expressing the stemness^67^ and proliferation^68^ markers (SOX2, Ki67) vs. the differentiation marker TH^38^ in undifferentiated cells, cells differentiated on well plate (2D) and cells differentiated in cryogels. c) Gene expression comparison for the same three groups of samples. Statistical analysis in b) and c) was carried out in 2 successive steps: First, the overall success of differentiation was assessed by comparison of the undifferentiated samples with the pooled differentiated samples (2D and cryogels). In a second step, possible differences in 2D vs. cryogel differentiation were assessed by direct comparison. Student’s t-test on per-experiment averages with Holm-Sidak correction for multiple comparisons, n=3 to 4 independent experiments (biological replicates) with 3-5 cryogels or wells (technical replicates per biological replicate) analysed per experiment and condition, *p ≤ 0.05, **p ≤ 0.01, ***p ≤ 0.001, ns = not significant. Brightness proportionally adjusted for visibility (Ki67, SOX2 images).

**Figure 5.**
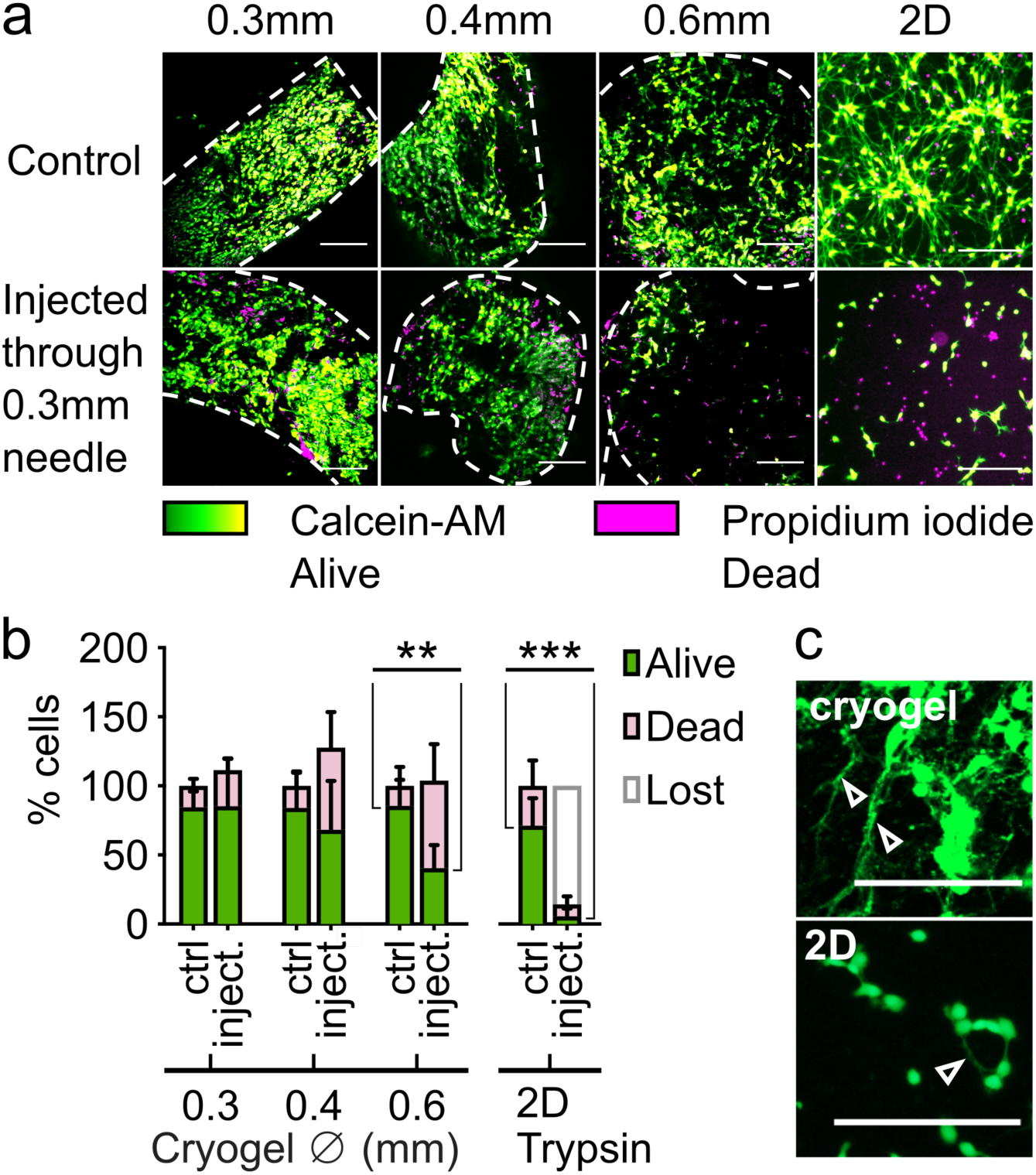
Effect of supporting Neurothread cryogel scaffold on viability of neurons upon passing through a 24 GA (0.3mm inner diameter) Hamilton syringe. a) Viability (calcein/PI) staining of cells, cultured on 3 diameters of Neurothread cryogels (0.3mm, 0.4mm and 0.6mm diameter, all 5mm length) or on a 2D well plate – either non-handled (control, ctrl) or 24 hours after passing through the 0.3mm inner diameter Hamilton syringe (in-vitro injection test). Image treatment to enhance visibility: Red (propidium iodide) replaced by magenta and linearly enhanced in brightness (2x), black-green-yellow lookup table applied to the calcein-AM fluorescence (indicated, associated brightness enhancement also 2x). b) Quantification of live and dead cells in control and injected samples for the 3 different diameter Neurothreads and the 2D culture reference. For each gel diameter, respectively the 2D condition, 100% represents the total amount of cells in the non-injected control. c) Neurites (white arrows) of cryogel-supported neurons or well 2D plate-cultured neurons after passing through the syringe. n=4-11. Student’s t-test with Holm-Sidak correction for multiple comparisons for a total of 4 comparisons of the recovery percentage of live cells between injected samples and associated control in b). **p ≤ 0.01, ***p ≤ 0.001, otherwise not significant. Scale bars represent 100 µm.

**Figure 6.**
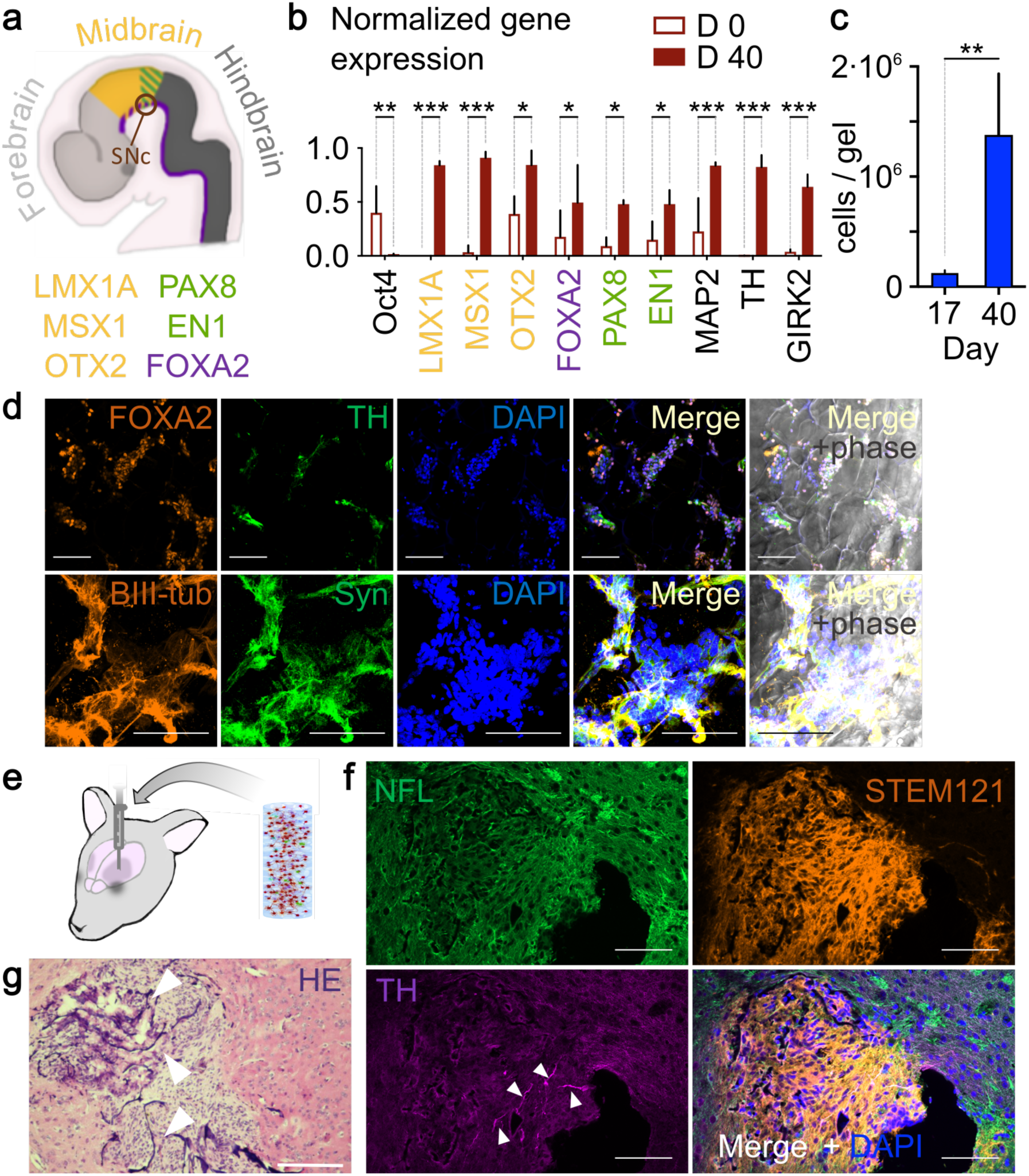
Differentiation and implantation of cryogel-supported hESC-derived D40 neurons. hESC-derived NPCs (day 16 or 17) were seeded on cryogels and differentiated until day 40 (D40). a) Scheme of midbrain marker expression around the substantia nigra pars compacta (SNc) area in the developing central nervous system^73^, with regional transcription factors.^38^ b) Midbrain fate gene expression profile of cryogel-differentiated neurons, assessed prior to initiation of differentiation (D0) and on day 40 (D40). Expression of markers of pluripotency – Oct 4^74^, ventral midbrain (LMX1A/FOXA2/MSX1)^38^, caudal midbrain (PAX8/EN1)^38^, rostral midbrain (OTX2)^38^, neurons (MAP2)^69^ and dopaminergic neurons (TH^38^, GIRK2, the latter more strongly expressed in the SNc^75^), relative to the house-keeping genes GAPDH/EEF1A1 and normalized to the highest expression observed in one of the technical replicates per gene. n=2-4 independent experiments, 2-4 cryogels per condition and experiment, statistical evaluation on the per-experiment averages, c) Cell numbers per gel at 24h after seeding on day 16 (i. e. D17) and day 40 (D40), estimated by Alamar Blue assay, n = 3-4 from single experiment, log transformation prior to statistical testing to account for exponential growth. d) Immunocytochemistry of FOXA2 and TH and d) BIII tubullin and Synapsin I in D40 hES-derived cells inside cryogels. e) D40 cryogel-supported neurons were injected into murine striatum. e) Hematoxylin and eosine (HE)-stained slice 4 weeks after graft injection. Arrows show examples of hematoxylin-stained cryogel walls, brightness proportionally increased for visibility. f) Immunohistochemistry of an adjacent slice, showing surviving human cells (STEM121), as well as neurofilament 200 (NFL)- and TH-positive cells and fibers in the area of the graft. Grey arrows show TH-positive cell bodies; TH-positive neurites are widely present in both host striatum and graft. Scale bars everywhere represent 100 µm. In d), Student’s t-test was used to compare conditions, followed by Holm-Sidak correction for multiple comparisons. *p ≤ 0.05, **p≤0.01, ***p≤0.001 otherwise not significant.

Dunnett’s^49^ or Tukey-Kramer’s^50^ multiple testing procedures were applied per subfigure if ANOVA was performed, otherwise the Holm-Sidak^51^ multiple testing procedure was used. The adjusted P-values are reported. Data tabulation and averaging was carried out in Excel, Version 14.4.2, and statistical analysis in Graphpad Prism version 8.3.0.

## Results

### Selection of adhesion molecules and optimization of scaffold functionalization

Our target is to enable *in-vivo* transplantation of mature dopaminergic neurons by the design of Neurothreads: cell-protective cryogel scaffolds for stereotactic injection. Our first step towards this goal was the synthesis and cell-adhesive functionalization of carboxymethylcellulose (CMC) cryogels.

We used a previously published protocol^20^ for the synthesis of the CMC cryogels. This yielded sponge-like structures (Fig. 1c), with known brain-like mechanical properties^20^. For neural adhesion to these sponge-like, extremely compressible scaffolds, we require efficient, yet highly biocompatible immobilization of neural adhesion molecules. We particularly wanted to avoid potentially toxic polycation components^52^ previously used with this system, such as PLO^20^, to avoid slow leakage^53^ after implantation. Hence, we investigated covalent immobilization of neural adhesion molecules by using a zero-length carbodiimide crosslinker, EDC^54^.

We optimized chemical conditions to immobilize three commonly used neural adhesion molecules – laminin 111, fibronectin and collagen IV^55,56^. For these experiments we used cryogel discs of 4 mm diameter and 1 mm height, allowing evaluation of protein immobilization by determination of the amino-group content with the amine-reactive dye rhodamine-isothiocyanate RITC^40^ in 96-well plate format (Fig. 2a). Our immobilization procedure consists of two separate steps, namely a protein adsorption step at controlled pH, followed by the reaction with EDC to achieve covalent, permanent immobilization (Fig. 2a). For the adsorption step, we investigated the use of a series of buffers between pH 4 and pH 8. We find increasing immobilization efficiency with decreasing pH for all three extracellular matrix (ECM) molecules tested (Fig. 2b, P=6.3*10^−6^, 4.5*10^−6^ and 7.5*10^−3^ for laminin, fibronectin and collagen IV respectively by F-tests for the explained vs. residual variance in non-linear fitting for inhibition of binding by increasing pH with a Hill coefficient of 1). The inflection points in the vicinity of the isoelectric points of the ECM proteins^57^ suggest that protein adsorption is strongly enhanced by electrostatic attraction between positively charged ECM molecules at lower pH and negatively charged carboxylgroups of the CMC scaffolds^58^. Indeed, in the most efficient conditions, a majority of the applied ECM molecules is immobilized. As a compromise between the requirement for efficient immobilization and usage of near physiological conditions to avoid protein denaturation, we used the pH 6 buffer for laminin and collagen IV adsorption and the pH 4 buffer for fibronectin adsorption in all further experiments.

The immobilization efficiency as quantified by RITC uptake shows a non-linear dependency on the crosslinker (EDC) concentration (Fig. 2c). The highest apparent protein binding is indeed obtained at intermediate EDC concentrations (3mg/mL), and a quadratic (P=0.01) rather than linear (P=0.29) regression appropriately fits the observed values. Of note, rather than quantifying protein binding directly, the uptake of the reactive dye RITC is dependent on available reactive groups such as amines and thiols.^40^ Chemical masking, by excessive amide bond formation^58^ with carboxyl groups from the carboxymethylcellulose and likely also by internal reaction between adjacent carboxyl- and aminogroups in the ECM proteins themselves^59^ probably explains this nonlinear behaviour.

Having established an efficient chemical protocol for covalent immobilization of neural adhesion molecules to the cryogel scaffolds, we next compared survival and neurite spread in the biofunctionalized cryogels (Fig. 3). For this, we seeded and neuronally differentiated the midbrain model cell line LUHMES^43^ in variously functionalized cryogel disks (Fig. 3a). In these experiments, we included not only negative, non-coated control cryogels, but also positive controls based on Matrigel. This commonly used strongly cell-adhesive ECM extract^60,61^ consists of a heterogeneous mix of adhesion proteins (wherein 60% laminin and 30% collagen IV), but also growth factors and many other components^62–64^.

We quantified neural spreading by the ratio of neurite objects to nuclei (eq. 1 in the Material and Methods section). In accordance with morphological observation (Fig. 3b), the highest neural spreading scores thus defined were observed in Matrigel- and laminin-functionalized gels (P=0.025 respectively P=0.0045 vs. uncoated control, unpaired t-test with Dunnetts correction after significant ANOVA). Fibronectin and collagen IV functionalization gave results similar to the negative control (P=0.83 respectively P=0.97, Fig. 3c). For collagen IV, the observed lack of cell adhesion might be at least in part linked to lower immobilization efficiency (Fig. 2b), while fibronectin on its own is apparently insufficient for neural spread in our conditions. The differences in neural spreading were primarily due to the cryogel functionalization and not viable cell numbers, which were similar for all conditions including controls (Fig. 3d, all P-values non-significant). This unanticipated finding of retention of similar cell numbers is explained by the formation of cell aggregates confined within the large pores of the cryogel, apparently permitting full maintenance of cell viability in the differentiating LUHMES cells. Further micromorphological analysis in Supplementary 1 (Figure S1) corroborates this: the number of nuclei is similar across the variously functionalized cryogels (Fig. S1a), only the neurite spreading varies as a function of cryogel modification (Fig. S1b).

Our results regarding neurite spreading are concurrent with previous studies showing that laminin promotes neurite spread and stimulates cell differentiation into the neuronal lineage^65,66^. Matrigel, while providing positive control results for neurite spread, would not be suitable for clinical applications, considering its origin and variable composition^62–64^. We thus used laminin 111 in the subsequent experiments.

In Fig. 2c, we found that the highest apparent protein immobilization efficiency was observed at intermediate, and not the highest EDC concentrations tested. Accordingly, we also investigated neurite spreading as a function of EDC concentration for laminin functionalization. We again observe a bell-shaped curve indicating an optimum EDC concentration. Among the values investigated, the best fit is indeed obtained with a parabolic response with a maximum at the 1mg/mL point (P=0.012, using the DAPI nuclei count as a covariate to reduce variability), whereas neither similar regression with a linear dependency on the EDC concentration (P=0.06) nor a parabolic dependency with a maximum imposed at 3mg/mL (P=0.89) gave significant results. The differences in neurite scoring are again due to cell morphology rather than cell numbers (Supplementary 1, Fig. S1c and S1d). Overall, 1mg/mL EDC most likely represents the best compromise between the need of covalent protein attachment to the scaffold to prevent protein leakage^37^ and persistence of unmodified side chains for interaction with cells. Hence, we use this concentration in all further experiments.

### Differentiation on cryogels vs tissue culture well plates

Having defined suitable cell adhesive conditions allowing for neurite spread, we next investigated whether the cryogel culture would influence the cell differentiation phenotype, or whether the 2D protocols can be transferred to our 3D cell culture system without change in differentiation efficiency.

To assess the effect of cryogel culture on dopaminergic differentiation, we seeded and differentiated the midbrain neural stem cell progenitor cell line LUHMES either on PLO/laminin-coated 2D well plates or in laminin-coated disk-shaped cryogels. Immunocytochemistry (Fig. 4a, Fig. 4b) showed successful differentiation in 2D and 3D conditions (significant decrease in the stemness and proliferation markers SOX2 and Ki67, P=0.036 respectively P=3.6*10^−8^, significant increase in dopaminergic marker tyrosine hydroxylase^38^ TH, P=0.0094, Holm-Sidak multiple testing correction). However, when comparing differentiation in the 3D cryogels vs. the 2D plates (Fig. 4b), there was no significant difference in cell numbers positive for either the stemness marker^67^ SOX2 (P=0.88), the proliferation marker^68^ Ki67 (P=0.97) or the catecholamine neuron marker^38^ TH (P=0.38). Analysis of gene expression of SOX2, Ki67, TH and MAP2 (microtubule-associated protein 2)^69^ showed similar results, with successful differentiation in both 2D and 3D conditions, but with no difference in gene expression between the the 3D and 2D differentiation conditions (Fig 4c). These results imply that neither the cryogel itself, nor the conformation of the cells acquired within it have any significant effect on the differentiation outcome.

### Injectability of Neurothread-supported neurons

With successful neural adhesion and differentiation protocols in place, we next proceeded with *in-vitro* injection tests with differentiated neurons. For this, we made once more use of the differentiation of LUHMES cells, seeded on cylindrical Neurothread cryogels with the aim of stereotactic minimally invasive delivery. In light of animal experiments in mice, we molded Neurothread cryogels with a cylindrical shape of 5mm length and a maximum diameter that would still preserve cell viability after handling and passing it through a 24 GA (0.311mm inner diameter according to the supplier’s notice) Hamilton syringe. In preliminary testing, we found 0.3mm diameter Neurothreads to be the smallest Neurothreads tolerant to handling, while Neurothreads of 0.6mm diameter were the largest possible to non-destructively fit into a the Hamilton needle, in agreement with previously described complete tolerance to up to 75% mechanical compression^20^. For assessment of neural viability during the injection and handling procedures, we therefore produced Neurothread cryogel carriers of three diameters: 0.3, 0.4 and 0.6 mm. For these Neurothreads, the passage through the Hamilton syringe implies 0%, 40%, respectively 70% cross-sectional compression.

We functionalized the Neurothread scaffolds with laminin, followed by seeding and differentiation of LUHMES cells for 7 days. Following this, part of the scaffolds of each size were left undisturbed, while another part was taken up into the syringe needle, retained for one minute and injected into a new well with fresh media. 24 hours later, injected and control gels of all sizes were labeled with calcein AM and propidium iodide (PI), to distinguish between live cells and nuclei of dead cells^70,71^. Well-plate differentiated LUHMES cells^43^ - either dissociated and passed through Hamilton or left in culture - were used as 2D reference.

Fig. 5a shows micrographs of live/dead stained Neurothread cryogels respectively 2D well plate cultures 24h after injectability testing. Visually, cell viability is largely conserved in the 0.3mm and 0.4mm cryogels as compared to their respective control, while the 0.6mm cryogels and 2D culture conditions shows substantial cell death or loss. Quantification of the recovery of viable and dead cells relative to non-injected control cell numbers in Fig. 5b confirms and nuances this. Statistical analysis of the number of viable cells relative to the respective non-injected controls shows a statistically significant two-fold decrease for the 0.6mm gels (P=0.0067), but no significant differences for the 0.3mm and 0.4mm gels (Fig. 5b). This implies some tolerance of the neural network to handling and compression, as reported previously^20^, but also defines a practical upper limit to 0.4mm diameter Neurothreads. In terms of recovery of life cells, all Neurothread cryogels perform much better than the dissociation of 2D cultures (P<0.0001 for the 0.3mm and 0.4mm gels, P=0.025 for the 0.6mm gels). Indeed, dissociation of the 2D cultures leads to a relative loss of viable cell numbers by more than an order of magnitude (P=5.4*10^−9^). This massive cell loss is associated with mechanical damage, as at 24h post-injection, at best short neurites could be observed in the dissociated cultures, whereas the extended cell morphology was better conserved on the cryogel matrices (Fig. 5c).

The results obtained prompted us to further investigate the mechanisms of loss of cell viability during the injection procedure. As shown in Supplementary Figure S2, the main effect of the cryogels is to protect the cells during handling, as for all three sizes, viable cell counts during the necessary gel handling steps (without passage through the Hamilton needle) is conserved. On the contrary, dissociation and replating of the 2D cultures fully accounts for the cell loss observed in Fig. 5b. Hence, the primary effect of the cryogels is to provide protection during mature neuron handling, with substantial, but nevertheless limited protection against compression.

Finally, to better anticipate the *in vivo* injection procedure, we injected 0.4mm diameter Neurothread cryogels, containing LUHMES-derived neurons, into an agar brain phantom^47^ (Supplementary Figure S3a). The cryogels were incubated inside the media-containing phantom for two hours – a timeframe estimated to allow the observation of most of the acute effect on the injected cells^72^, yet not long enough for possible medium depletion from the phantom. No cell viability decrease in the cryogel-seeded cells was observed (Supplementary Figure S3b), further supporting the choice of the 0.4mm Neurothreads for *in-vivo* experiments.

### hESC differentiation and in-vivo injection

As a next step, we sought to transfer an established protocol for the production of human dopaminergic neurons from embryonic stem cells to our cryogel system. We use the protocol by Nolbrant et al.^38^, as it is potentially compatible with Good Manufacturing Practice and clinical application^38^. For the assessment of differentiation efficiency, we pre-differentiated the human ESC line HS420 for 16-17 days and seeded the resulting neural progenitor cells onto laminin111-coated, disk-shaped cryogels for terminal differentiation.

Fig. 6b shows the change of gene expression levels occurring from the hESC state (D0) to the final neural state on the cryogels (D40), with the most important regionalization markers being outlined in Fig. 6a. We observed significant loss of expression of the pluripotency marker Oct4^74^, and a significant increase in transcription factors associated with midbrain fate (LMX1A, MSX1, FOXA2, OTX2)^38^ and caudalization (EN1, PAX8)^38^. In addition, the neural maturity marker MAP2^69^, as well as the dopaminergic fate markers (TH^38^ and GIRK2^75^) also exhibited the anticipated significant increase. This marker expression profile is in line with literature^38^, confirming the compability of cryogel culture with an established differentiation protocol^38^ also in the context of hESC differentiation. By immunocytechemistry (Fig. 6d), we find FOXA2 and TH-positive cells (8.26 ± 3.38 % TH^+^ and 7.94 ± 3.99 % TH^+^/FOXA2^+^) in a comparable amount to the published results^38^. BIII-tubullin-positive neurites and Synapsyn-1 expression was also observed (Fig. 6d). The widespread presence of the latter indicates that indeed, we have obtained relatively mature, histologically connected neurons as the Synapsin-1 protein is part of the mature synaptic machinery^76–78^. In terms of cell amount, we observed a major increase from day 17 (24h after seeding) to day 40 (Fig. 6c, P=0.0031, t-test after log transform), showing that NPCs continue to proliferate before acquiring a mature phenotype.

Having successfully established hESC differentiation on our cryogel system, we next produced Neurothread cryogels (0.4mm diameter) for a pilot experiment with stereotactic injection of cryogel-supported hESC-derived neurons into the striatum of immunosuppressed mice (Fig. 6e). One month after the injection, H&E staining of striatal slices showed the well-preserved structure of the implanted cryogel (Fig. 6g) and a large number of nuclei residing within it’s pores. Immunofluorescence staining of sequential slices showed high density of compact tissue of human origin as identified by the human cytosolic marker^79^ STEM121 (Fig. 6f). Immunohistochemistry against neurofilaments supported a neural phenotype of the transplanted cells. Tyrosine hydroxylase (TH) labelling revealed a dense network of TH-positive neurites, and occasional TH positive cell bodies.

This concludes the proof of concept of the protective effect of macroporous cryogels during the transplantation of a histologically mature neural network during intra-cerebral stereotactic transplantation procedures.

## Discussion

Our aim is to provide a Neurothread cryogel delivery system for intracerebral implantation of mature neurons derived *in vitro.* We expect this approach to open the possibility to enhance neuronal maturation and ensure appropriate midbrain dopaminergic differentiation of the graft, providing an opportunity to increase efficacy of restoration of motor function and decrease the risk of devastating side-effects, like tumorigenesis and GIDs, currently associated with stem cell therapy in PD^4^. Furthermore, this platform can potentially be used for other pathologies stemming from focal neuronal loss, including other focalized neurodegenerative diseases such as Huntington’s disease and stroke.^80–82^

The use of porous, rather than physically encapsulating, scaffolds requires implementation of adhesion motives for cell binding^18,20,25^. To achieve effective carbodiimide-mediated protein immobilization^40^, we use a distinct adsorption step below the isoelectric point of the ECM molecules. Interestingly, this allows us to circumvent the otherwise cumbersome need for peptide coupling enhancers such as N-hydroxysuccinimide^83^. Possibly, the spatial proximity induced by the electrostatic interaction allows to prevent hydrolysis of the activated esters^40^ prior to amine-coupling. Ultimately, the efficient covalent functionalization avoids the use of leachable and potentially toxic polycations such as PLO^20^. The result also demonstrates that compared to previous studies^20^, and also typical 2D neural cultures and neural differentiation^79^, that the polycation coating might be dispensable provided alternate efficient means of immobilization.

Regarding neural adhesion, our results indicate a positive effect of laminin 111 on neurite elongation, in agreement with known results^65,66^. We indeed find the protein to promote neurite elongation in an extent similar to Matrigel when used in similar concentrations. Whereas using Matrigel for clinical applications would be difficult^84^, the sarcoma-extracted^63^ laminin 111 used here could be replaced with a recombinant version in the next steps. This would ensure substantial cell adhesion, yet provide a potential pathway to clinical studies.

We show successful differentiation of immortalized mesencephalic neural precursors, LUHMES cells^43^, with similar expression of pluripotency, proliferation and differentiation markers within the cryogels and on culture plates. Likewise, upon differentiation of hESCs on cryogels, we detect the anticipated gene expression patterns^55^ with an increase in a series of typical midbrain, neuronal and catecholamine neuron markers, indicating successful differentiation of these hPSCs into ventral mDA neuron phenotype on the cryogels. There seems to be little influence from the cryogel geometry on the differentiation efficiency. We attribute this to the biologically inert nature of the CMC backbone used for scaffold synthesis^85^. Indeed, while we find the CMC to enable facile modification through carbodiimide chemistry, this study confirms that this molecule provides little adhesion or other biological signals to cells by itself. We therefore anticipate various emerging cell sources and protocols to be easy to adopt for the use with our biomaterial, broadening the scope of applications and increasing the efficacy of cell therapy in both Parkinson’s and other diseases.

Our *in-vitro* injectability tests with differentiated LUHMES cells confirm a remarkable capacity of the cryogel scaffolds to protect fragile cells with extended neurites.^20^ There was indeed no significant loss of viable cells during handling and the moderate 40% compression required for stereotactic delivery of 0.4mm diameter Neurothreads through 0.3mm inner diameter Hamilton needles for stereotactic delivery. In stark contrast, more than 90% of viable cells were lost when dissociating differentiated 2D cultures. The *in-vivo* results obtained here provide further proof-of-principle of efficient transplantation of very high-density cell assemblies, with formation of a new neural tissue *in situ*. The compression of up to 40% without substantial cell loss additionally permits minimally invasive injection of a neuron-filled scaffolds somewhat larger than the needle track, capable of gentle swelling to the available tissue space. This minimizes the necessary needle diameter and thus risk of complications during surgery.

We observed a significant increase in the amount of cells during maturation from an NPC to a mature neuronal state (Fig, 6c). Although further analysis at longer time-points is certainly required, we consider this result as an *in-vitro* confirmation of the potential risk of *in-vivo* tumorigenesis, one of the main challenges associated with transplantation NPCs^86,87^. Our results highlight the need for tight control of proliferation and terminal differentiation. This underscores the importance of emerging approaches such as the incorporation of a suicide gene for proliferating cells into the graft^86^, selection of a suitable progenitor population, expressing specific surface ventral midbrain markers^88^, or by trans-differentiation of more mature cells^89^. It also underpins the importance of approaches such as the one investigated here that enable prolonged maturation prior to implantation, offering the possibility of later quality control points.

During a pilot injection of cryogel-supported hESC-derived D40 neurons, we observed preservation of these mature neurons within the scaffold – including cells positive for TH - one month after the grafting procedure. This provides proof-of-concept for transplantation of a relatively mature network of neurons. This achievement provides a unique opportunity to test and verify the strategies aimed at ensuring proliferation-free, terminal differentiation. The possibility to transplant neurons with mature morphology and synaptic connectivity in a fully porous scaffold also offers fundamentally novel possibilities in the emerging field of pre-assembled neural circuitry. Some initial steps in this direction have indeed been made with tubular micro-tissue engineered neural tracts^28,29^ or chemical definition of reconstruction pathways.^90^ But the difficulty to maintain neural viability, yet enable rapid interaction with the surrounding tissue during transplantation has so far let this field largely unexplored^19^.

## Conclusions

The versatility offered by a biologically inert, yet nearly arbitrarily modifiable scaffold enabling mature neuronal transplantation lets us anticipate new perspectives regarding dopaminergic neural transplantation for the treatment of Parkinson’s disease. The Neurothread platform enables extensive *in-vitro* culture, differentiation and characterization prior to efficient *in-vivo* transfer of preformed neural networks and has the potential to powerfully leverage neural maturation and differentiation quality control. Beyond Parkinson’s disease, the Neurothread cryogel platform developed here may offer a building block in the emerging transplantation of intact neural circuitry, with the potential to offer a bridge between the rapidly developing field of organoid and engineered tissue culture and neural transplantation with clinical application.

## Acknowledgments

We would like to thank Patrick Burch from Volumina-Medical SA for his help with cryogel premix formulations; Benoît Desbiolles from École Polytechnique Fédérale de Lausanne (EPFL) for assistance with electron microscopy imaging; and Antoine Marteyn, Manon Locatelli and Vannary Tieng from the University of Geneva (UNIGE) for guidance regarding stem cell differentiation procedures, and Vincent Jaquet and Joé Bréfie-Guth (UNIGE) for assistance in animal experiments. We further thank the personnel at the following core facilities for their kind help with training, setup and technical challenges: the Bioimaging Core Facility, the Genomics Core facility, the READS unit and the Histology facility of the Faculty of Medicine of the University of Geneva, as well as the Bioelectronmicroscopy core facility of the EPFL. We also specially thank the caretakers of the UNIGE animal facilities for their maintenance and support. Finally, we would like to acknowledge funding sources from the Gebert-Rüf foundation (GRS-043/15 to A.B.) and the Swiss National Science Foundation PP00P2_163684 and PZ00P2_161347 to T.B.).

## Supplementary Information List (separate file)

- Supplementary 1: Neurite and nuclei counts for optimization of neural spread
- Supplementary 2: Neural cell survival during handling
- Supplementary 3: Injection into a brain phantom
- Supplementary 4: List of antibodies
- Supplementary 5: List of primers

## Author contributions

T.B. and A.F. initiated the project. A.F., K.H.K., A.B. and T.B. designed the study. A.F. and F.B. developed the biomaterial coating. A.F. performed the *in-vitro* experiments with the advice and help of L.E. and O.P. A.F. assessed injectability and peformed *in-vivo* experiments with the advice of K.H.-K., A.B. and T.B. A.F. and T.B. analysed and curated the data, and compiled the figures. A.F. and T.B. wrote the primary manuscript, all authors thoroughly reviewed the manuscript, and approved of the final form of the manuscript.

## Data availability

The partially processed data required to reproduce these findings are available to download from https://doi.org/10.5281/zenodo.3608207

